# Sequence dependence of biomolecular phase separation

**DOI:** 10.1101/2020.09.24.312330

**Authors:** Benjamin G. Weiner, Yigal Meir, Ned S. Wingreen

## Abstract

Cells are home to a wide variety of biomolecular condensates - phase-separated droplets that lack a membrane. In addition to nonspecific interactions, phase separation depends on specific binding motifs between constituent molecules. Nevertheless, few rules have been established on how these specific, heterotypic interactions drive phase separation. Using lattice-polymer simulations and mean-field theory, we show that the sequence of binding motifs strongly affects a polymer’s ability to phase separate, influencing both phase boundaries and condensate properties (e.g. viscosity and polymer diffusion). We find that sequences with large blocks of a single motif typically form more inter-polymer bonds which promote phase separation. Notably, the sequence of binding motifs influences phase separation primarily by determining the conformational entropy of self-bonding by single polymers. This contrasts with systems where the molecular architecture primarily affects the energy of the dense phase, providing a new entropy-based mechanism for the biological control of phase separation.

## Introduction

Understanding how biological systems self-organize across spatial scales is one of the most pressing questions in the physics of living matter. It has recently been established that eukaryotic cells use phase-separated biomolecular condensates to organize a variety of intracellular processes ranging from ribosome assembly and metabolism to signaling and stress response (***Hyman et al., 2014; Banani et al., 2017; Boeynaems et al., 2018***). Biomolecular condensates are also thought to play a key role in physically organizing the genome and regulating gene activity (***Hnisz et al., 2017; Sabari et al., 2018; Shin et al., 2018***). How do the properties of these condensates emerge from their components, and how do cells regulate condensate formation and function? Unlike the droplets of simple molecules or homopolymers, intracellular condensates are typically composed of hundreds of molecular species, each with multiple interaction motifs. These interaction motifs can include folded domains, such as in the nephrin-Nck-N-WASP system for actin regulation (***Li et al., 2012***), or individual amino acids in proteins with large intrinsically disordered regions (IDRs), such as the germ granule protein Ddx4 (***Nott et al., 2015***). While the precise sequences of these motifs are believed to play a major role in determining condensates’ phase diagrams and material properties, the nature of this relation has only begun to be explored (***Brangwynne et al., 2015; Alberti et al., 2019; Hicks et al., 2020***). As a result, it remains difficult to predict the formation, properties, and composition of these diverse functional compartments.

Previous studies have established important principles relating phase separation to the sequence of nonspecific interaction domains such as hydrophobic or electrostatic motifs (***Lin et al., 2016; Das et al., 2018; McCarty et al., 2019; Statt et al., 2020***). However, in many cases condensate formation and function depend on specific interactions which are one-to-one and saturating (***Banani et al., 2017***). These can include residue-residue bonds, bonds between protein domains, protein-RNA bonds, and RNA-RNA bonds. Such one-to-one interactions between heterotypic domains are ubiquitous in biology, and recent studies have enumerated a large number of examples in both one-component (***Wang et al., 2018***) and two-component (***Ditlev et al., 2018; Xu et al., 2020***) systems (e.g. cation-pi bonds between tyrosine and arginine in FUS-family proteins, bonds between protein domains in the SIM-SUMO system). Another important example is RNA phase separation in “repeat-expansion disorders” such as Huntington’s disease and ALS. There, phase separation is driven by specific interactions between nucleotides arranged into regular repeating domains, and it has recently been shown that the repeated sequence pattern is necessary for aggregate formation (***Jain and Vale, 2017***). In spite of the biological importance of such specific interactions, their statistical mechanical description remains undeveloped. Here, we address the important question: what is the role played by sequence when specific, heterotypic interactions are the dominant drivers of phase separation?

Specifically, we analyzed a model of polymers with specific, heterotypic interaction motifs using Monte Carlo simulations and mean-field theory. We found that motif sequence determines both the size of the two-phase region and dense-phase properties such as viscosity and polymer extension. Importantly, sequence acts primarily by controlling the entropy of self-bonds. This suggests a new paradigm for biological control of intracellular phase separation: when bonds are specific and saturating, the entropy of *intra*molecular interactions can be just as relevant as the energy of *Inter*molecular interactions.

## Results

How does a polymer’s sequence of interaction motifs affect its ability to phase separate? To address this question, we developed an FCC lattice model where each polymer consists of a sequence of “A” and “B” motifs which form specific, saturating bonds of energy *ε* (Fig. 1 (a) and 1 (b)). Monomers on adjacent lattice sites also have nonspecific interaction energy *J*. For each sequence, we determined the phase diagram, which describes the temperatures and polymer concentrations at which droplets form. To enable full characterization of the phase diagram including the critical point, we used Monte Carlo simulations in the Grand Canonical Ensemble (GCE): the 3D conformations of the polymers are updated using a predefined move-set, and polymers are inserted/deleted with chemical potential *μ*. (See Methods and Materials for details.) For each sequence, we determined the critical point (temperature *T*_c_ and chemical potential *μ*_c_). Then for each *T* <*T*_c_ we located the phase boundary, defined by the value *μ*\* forwhich the dilute and dense phases have equal thermodynamicweight. Around this value of *μ*, the system transitions back and forth between the two phases throughout the simulation, leading to a polymer number distribution *P*(*N*) that has two peaks with equal weights (Fig. 1(c)) (***Panagiotopoulos et al., 1998***). The dilute and dense phase concentrations *ϕ*_dilute_ and *ϕ*_dense_ are the means of these two peaks. Multicanonical sampling was employed to adequately sample transitions (Methods and Materials).

**Figure 1.**
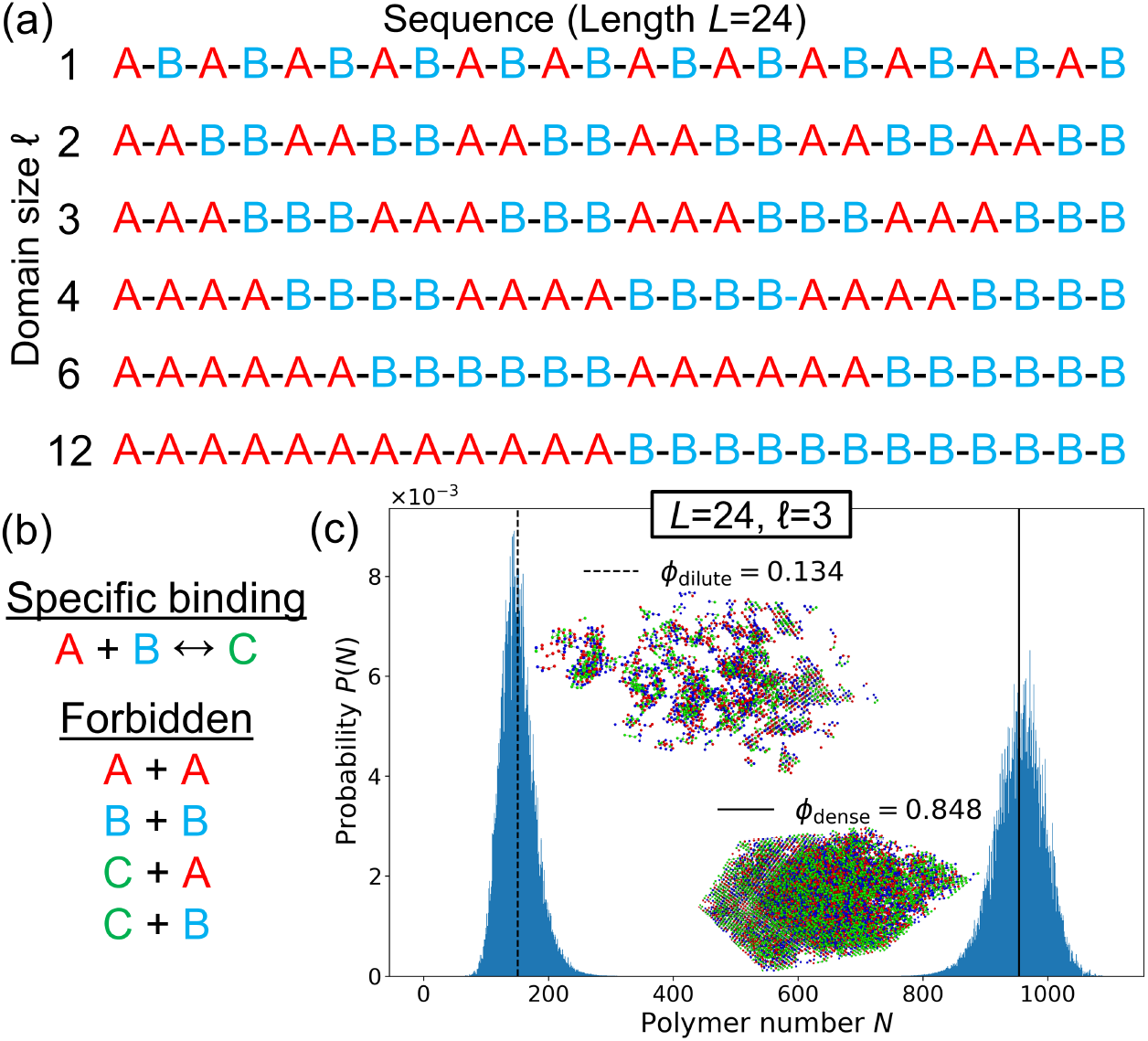
Lattice model for phase separation by polymers with one-to-one interacting motifs. (a) Each polymer is defined by its sequence of motifs, which come in types “A” (red) and “B” (blue). The class of sequences shown consists of repeated domains of As and Bs, labeled by their domain size *ℓ*. (b) In lattice simulations, an A and a B motif on the same lattice site form a specific, saturating bond (green) with binding energy *ε*. Monomers of any type on adjacent lattice sites have an attractive nonspecific interaction energy *J* = 0.05*ε*. A-A and B-B overlaps are forbidden. (c) Polymer number distribution *P*(*N*) at the phase boundary of the *ℓ* = 3 sequence (*βε* = 0.9287, *μ* = −9.9225*ε*). At fixed *μ* the system fluctuates between two phases. *Inset*: Snapshots of the GCE (fixed *μ*) simulation at *ϕ*_dilute_ and *ϕ*_dense_.

We first constructed phase diagrams for polymers with the six sequences shown in Fig. 1 (a), all with *L* = 24 motifs arranged in repeating domains, and all with equal numbers of A motifs and B motifs (*a* = *b* = 12 where *a* and *b* are the numbers of A and B motifs in a sequence). Each simulation contains polymers of a single sequence, and the sequences differ only in their domain sizes *ℓ*. Figure 2(a) shows the resulting phase diagrams, which differ dramatically by domain size, e.g. the *T*_c_ values for *ℓ* = 2 and *ℓ* = 12 differ by 20%. The absolute magnitude of the effect depends on the interaction energy scale *ε*, but we note that if the *T*_c_ for *ℓ* = 12 were in the physiological range around 300K, the corresponding 60K difference would render the condensed phase of *ℓ* = 2 inaccessible in most biological contexts. Despite this wide variation, Fig. 2(b) shows that rescaling by *T*_c_ and *ϕ*_c_ causes the curves to collapse. This is expected near the critical point, where all sequences share the behavior of the 3D Ising universality class (***Panagiotopoulos et al., 1998***), but the continued nearly exact data collapse indicates that (*T*_c_, *ϕ*_c_) fully captures the sequence-dependence of the phase diagram.

**Figure 2.**
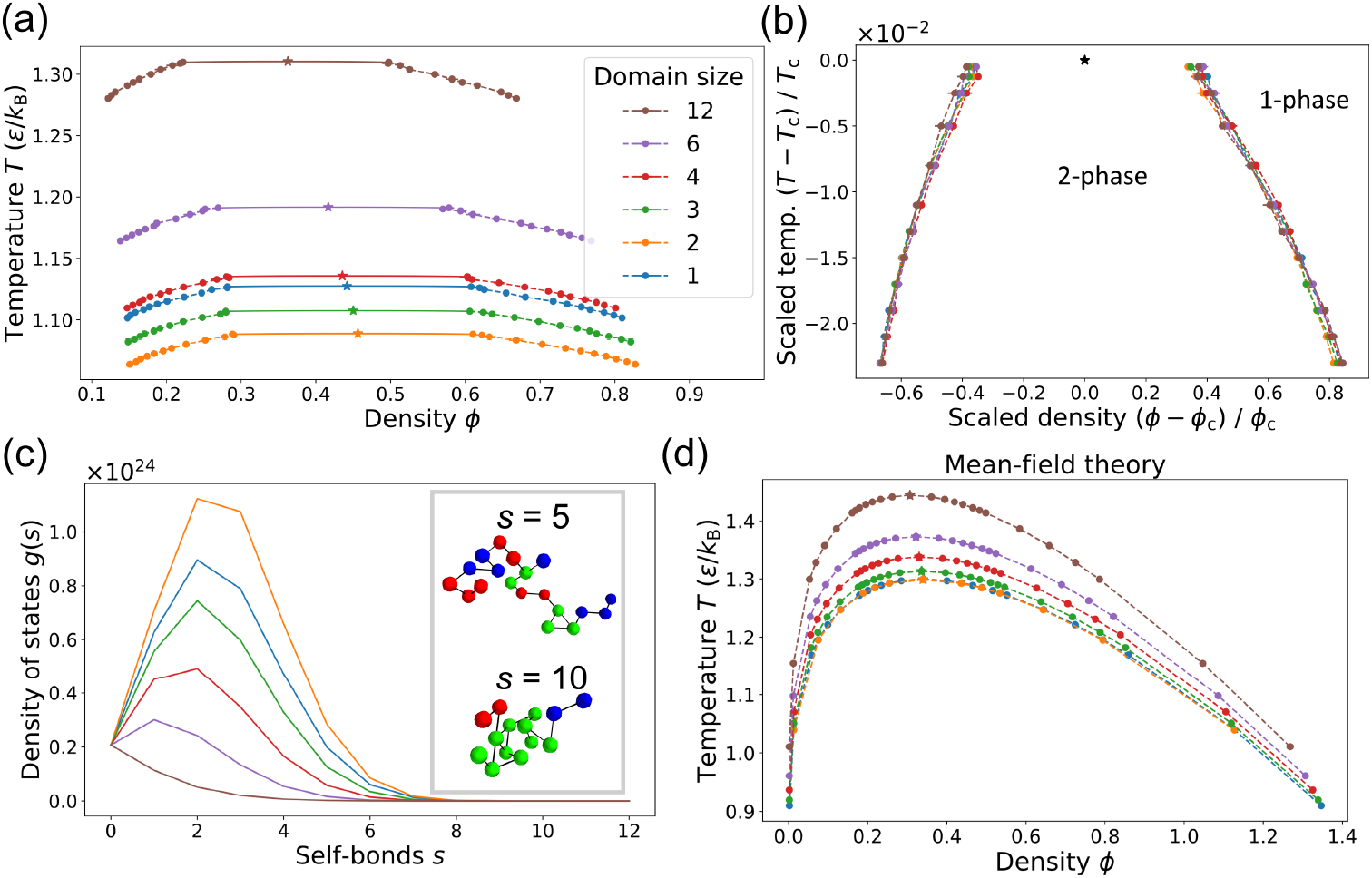
The sequence of binding motifs strongly affects a polymer’s ability to phase separate. (a) Binodal curves defining the two-phase region for the six sequences of length *L* = 24 shown in Fig. 1 (a). Stars indicate the critical points and the solid curves are fits to scaling relations for the 3D Ising universality class. Mean ± SD for three replicates. (Uncertainties are too small to see for most points.) Color key applies to all panels. (b) When rescaled by the critical temperature *T*_c_ and critical density *ϕ*_c_, the phase boundaries in (a) collapse, even far from the critical point. (c) The tendency to phase separate is inversely related to the density of states *g*(*s*), i.e. the number of ways a given sequence can form *s* bonds with itself. Inset: Snapshots of *ℓ* = 3 polymer with *s* = 5 (top) and *s* = 10 (bottom). Black lines show the polymer backbone. (d) Phase boundaries from mean-field theory using *g*(*s*) (Eq. 1).

Why does the sequence of binding motifs have such a strong effect on phase separation? Importantly, sequence determines the entropy of intra-polymer bonds, i.e. the facility of a polymer to form bonds with itself. This is quantified by the single-polymer density of states *g*(*s*): for each sequence, *g*(*s*) counts the number of 3D conformations with *s* self-bonds. For short polymers, *g*(*s*) can be enumerated, whereas for a longer polymers, it can be extracted from Monte Carlo simulations. Figure 2(c) shows *g*(*s*) for each of the domain sequences, obtained from Monte Carlo simulations. Sequences with small domain sizes have many more conformations available to them at all values of *s*. Intuitively, a sequence like *ℓ* = 2 allows a polymer to make many local bonds, whereas a sequence like *ℓ* = 12 cannot form multiple bonds without folding up globally like a hairpin. Such hairpin states are thermodynamically unfavorable at these temperatures due to the low conformational entropy, so it is more favorable for polymers like *ℓ* = 12 to phase separate and form trans-bonds with others, leading to a high *T*_c_ value. Even when *T* < *T*_c_ so that low-energy states with many bonds are favored, large-domain sequences have large two-phase regions because *g* (*s*) is small for all *s*. Thus, polymers with large domains form condensates over a much wider range of temperatures and concentrations.

This intuition can be captured by a simple mean-field theory that incorporates only single-polymer properties, namely *g*(*s*) and the number of A and B motifs per polymer, *a* and *b*. We calculate the free energy density of a state where each polymer forms *s* self-bonds and *t* transbonds (bonds with other polymers). We make two mean-field simplifications: 1) every polymer has the mean number of trans-bonds 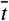, and 2) each polymer interacts with the others through a mean-field background of independent motifs. In contrast, the self-interaction is described by the full density of states *g*(*s*) extracted from single-polymer simulations. This leads to the following free energy density (see Appendix 1 for derivation):

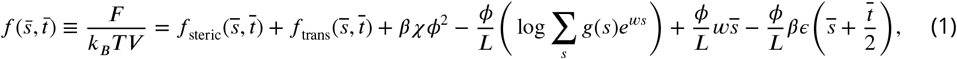

where *V* is the number of lattice sites, *χ* is the nonspecific-interaction parameter,

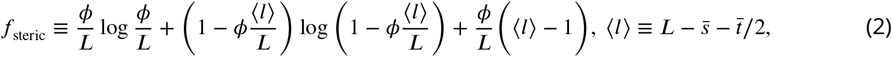

and

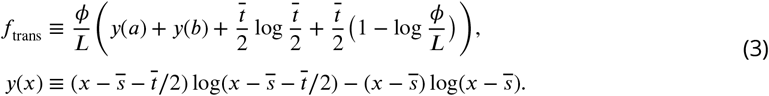

*f*_steric_ is the translational contribution from the number of ways to place polymers without overlap and *f*_trans_ is the entropy of forming 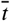 trans-bonds given 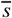 self-bonds, derived from the combinatorics of pairing independent motifs. The fourth term in Eq. 1 accounts for the self-bonding entropy, where *w* is the self-bond weight chosen to self-consistently enforce 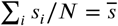. The next term is the Legendre transform compensating for *w*. (This allows us to estimate the entropy of 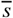 without assuming that 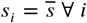. The procedure is akin to introducing a “chemical potential” *w* which fixes the mean number of self-bonds.) In the thermodynamic limit the partition function is dominated by the largest term, so we minimize Eq. 1 with respect to 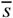 and 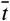 at each *ϕ* to yield *f* (*ϕ*) and determine the phase diagram.

Figure 2(d) shows the mean-field phase diagrams. In spite of the theory’s approximations, it captures the main patterns observed in the full simulations. Specifically, sequences with larger motif domains have larger two-phase regions and these extend to higher temperatures. (The mean-field *T*_c_ values differ from the simulations, but these could be tuned by the nonspecific-interaction parameter *χ*. Density fluctuations make it difficult to map *χ* to *J*, so we use the mean-field relation *χ* = *-VJz*/2 for simplicity.) Rescaling by *T*_c_ and *ϕ*_c_ also causes the mean-field phase boundaries to collapse (Appendix 4). Intriguingly, the mean-field theory does not correctly place the *ℓ* = 1 sequence in the *T*_c_ hierarchy. The single-polymer density of states *g*(*s*) suggests that *ℓ* = 1 should be similar to *ℓ* = 2, but its *T*_c_ is closer to to *ℓ* = 4. We trace this discrepancy to trans-bond correlations in the dense phase: the *ℓ* = 1 sequence tends to form segments of multiple bonds rather than independent bonds (see Appendix 2 for details). Overall, the success of the theory demonstrates that motif sequence mainly governs phase separation through the entropy of self-interactions. We capture this dependence, as well as corrections due to dense-phase correlations, in a simple “condensation parameter” described below.

Do these conclusions still hold if the motifs are not arranged in regular domains, and how do polymer length and motif stoichiometry affect phase separation? To address these questions, we located the critical points for three new types of sequences: 1) Length *L* = 24 sequences with *a* = *b* =12 in scrambled order, 2) domain sequences with *L* ≠ 24, and 3) sequences with *L* = 24 but *a* ≠ *b*. Each simulation contains only polymers of a single sequence. We find that the *T*_c_ hierarchy with respect to domain size *ℓ* is preserved across sequence lengths, so domain size is a robust predictor of phase separation (Appendix 4, Fig. 12). Figure 3(a) shows *T*_c_ and *ϕ*_c_ for the scrambled *L* = 24 sequences and for domain sequences of various lengths. *T*_c_ and *ϕ*_c_ are negatively correlated across all sequences because for low-*T*_c_ sequences, trans-bonds - and consequently, phase separation - only become favorable at higher polymer density.

**Figure 3.**
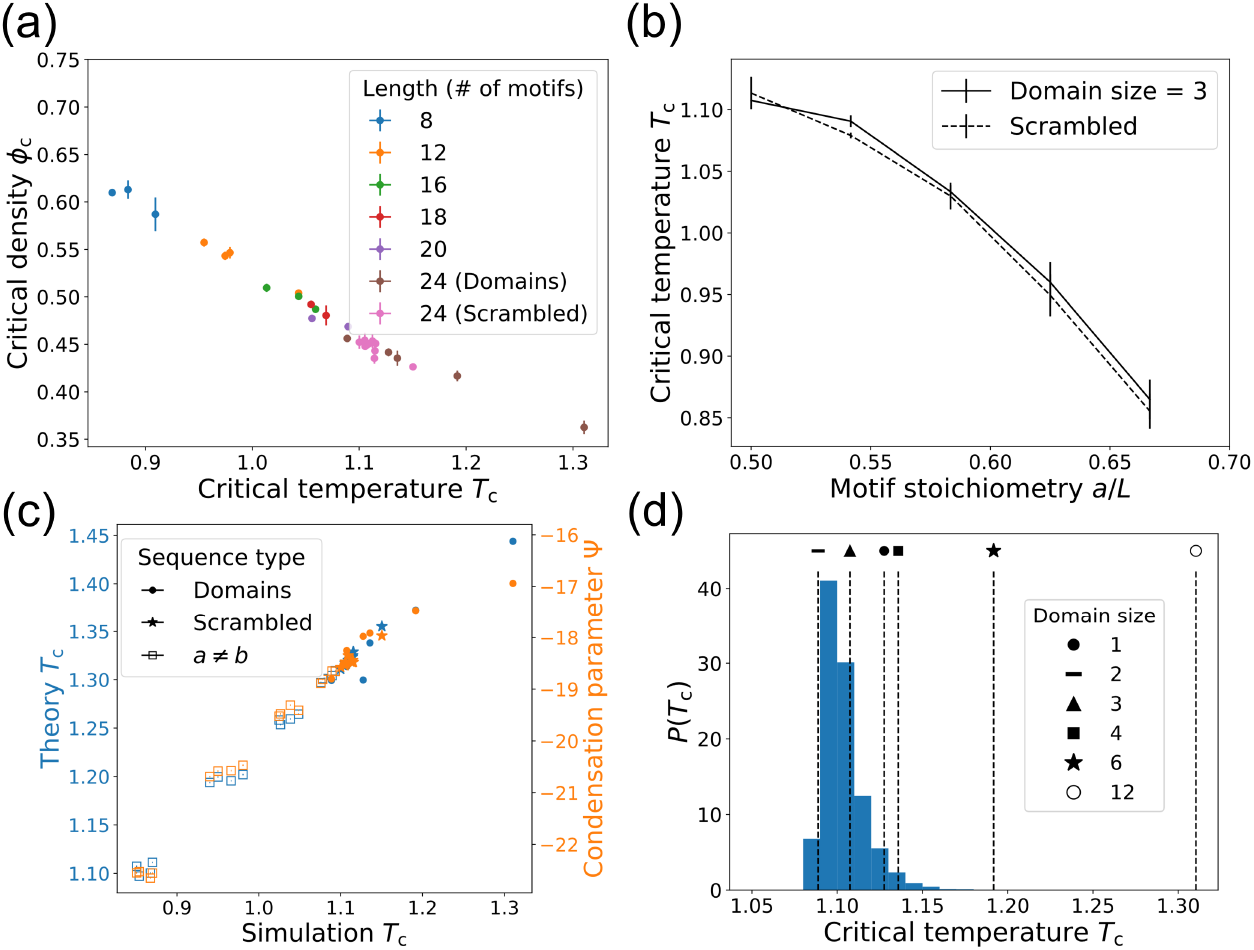
Ability to phase separate is determined by the sequence of binding motifs for polymers of different lengths, patterns, and motif stoichiometries. (a) *T*_c_ and *ϕ*_c_ for *L* = 24 polymers with scrambled sequences and domain sequences of various lengths. Mean ± SD over three replicates. (Temperature uncertainties are too small to see in (a) and (c).) (b) *T*_c_ as a function of motif stoichiometry *a/L*. The solid curve corresponds to *ℓ* = 3 sequences where a number of B motifs are randomly mutated to A motifs, and the dashed curve shows scrambled sequences. Mean ± SD over four different sequences. (c) *T*_c_ from Monte Carlo simulations versus mean-field theory (blue) and condensation parameter (orange) for domain sequences, scrambled sequences, and sequences with unequal motif stoichiometry, all *L* = 24. Mean ± SD over three replicates for simulation *T*_c_. (d) Distribution of *T*_c_ values for 20,000 random sequences of length *L* = 24 with *a* = *b*, calculated from Ψ values and the linear *T*_c_ versus Ψ relation for domain sequences. Domain sequence *T*_c_ values are marked.

The dashed curve in Fig. 3(b) shows *T_c_* for scrambled sequences with unequal motif stoichiometry. *T_c_* decreases as the motif imbalance grows because the dense phase is crowded with unbonded motifs, making phase separation less favorable. How does this crowding effect interplay with the previously observed effect of *g*(*s*)? Scrambled sequences are clustered near the *ℓ* = 3 sequence in (*T*_c_, *ϕ*_c_) space (Appendix 4, Fig. 11), so we generated sequences by starting with *ℓ* = 3 and randomly mutating B motifs into A motifs (Fig. 3(b), solid curve). The *ℓ* = 3 mutants follow the same pattern as the scrambled sequences, indicating that self-bond entropy and stoichiometry are nearly independent inputs to *T*_c_. This arises because motif flips have a weak effect on *g*(*s*) but a strong effect on dense phase crowding, giving cells two independent ways to control condensate formation through sequence.

The mean-field theory of Eq. 1 also captures the behavior of these more general sequences, as shown in Fig. 3(c). The critical temperatures from theory (blue markers) correlate linearly with the simulation *T*_c_ values. (The magnitude differs, but this is tuned by the strength of nonspecific interactions.) This agreement reinforces the picture that *T*_c_ is mainly governed by the relative entropy of intra- and inter-polymer interactions. The former is captured by *g*(*s*) and the latter depends on the motif stoichiometry. To capture these effects in a single number, we propose a condensation parameter Ψ which correlates with a sequence’s ability to phase separate (see Appendix 3 for a heuristic derivation):

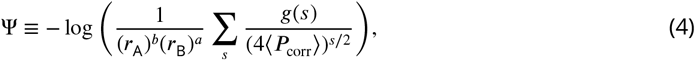

where *r*_A_ = *a/L* is the fraction of motifs that are A (and likewise for *r*_B_) and 〈*P*_corr_〉 is a simple metric for trans-bond correlations (See Appendix 2). A sequence with large Ψ has a high *T*_c_ because the dense phase is relatively favorable due to low self-bonding entropy, strong dense-phase correlations, or balanced motif stoichiometry. As shown in Fig. 3(c) (orange markers), this accurately captures the phase separation hierarchy of *T*_c_, including the correlation-enhanced *T*_c_ of the *ℓ* = 1 sequence.

Are domain sequences special? The space of possible sequences is much larger than can be explored via Monte Carlo simulations. However, we can use the condensation parameter to estimate *T*_c_ for any sequence without additional simulations. First, we estimate *g*(*s*) analytically and use this to approximate Ψ for new sequences. Then we use a linear fit of Ψ to the known *T*_c_ values for the domain sequences to estimate the critical temperature (details in Appendix 3). Figure 3(d) shows the distribution of critical temperatures calculated in this way for 20,000 random sequences with *a* = *b* = 12. Strikingly, the distribution is sharply peaked at low *T*_c_, similar to the domain sequences with *ℓ* = 2 or *ℓ* = 3. If particular condensates with high *T*_c_ are biologically beneficial, then evolution or regulation could play an important role in generating atypical sequences like *ℓ* = 12 with large two-phase regions.

The sequence of specific-interaction motifs influences not only the formation of droplets, but also their physical properties and biological function. Figure 4(a) shows the number of self-bonds in the dense phase relative to scaled temperature |*T* - *T*_c_|/*T*_c_. Density fluctuates in the GCE, so each point is averaged over configurations with *ϕ* within 0.01 of the phase boundary, and this density is indicated via the marker color (marker legend in 4(c)). The sequence ordering of self-bonds in the dense phase matches the sequence ordering of the single-polymer *g*(*s*), indicating that sequence controls intrapolymer interactions even in the condensate. Figure 4(b) shows the number of trans-bonds in the dense phase, plotted as in (a). Larger domains lead to more trans-bonds, even though the droplets are less dense. As temperature is reduced - and thus density is increased - the number of trans-bonds increases. Interestingly, even though the phase boundaries collapse to the same curve (Fig. 2(b)), different sequences lead to droplets with very different internal structures.

**Figure 4.**
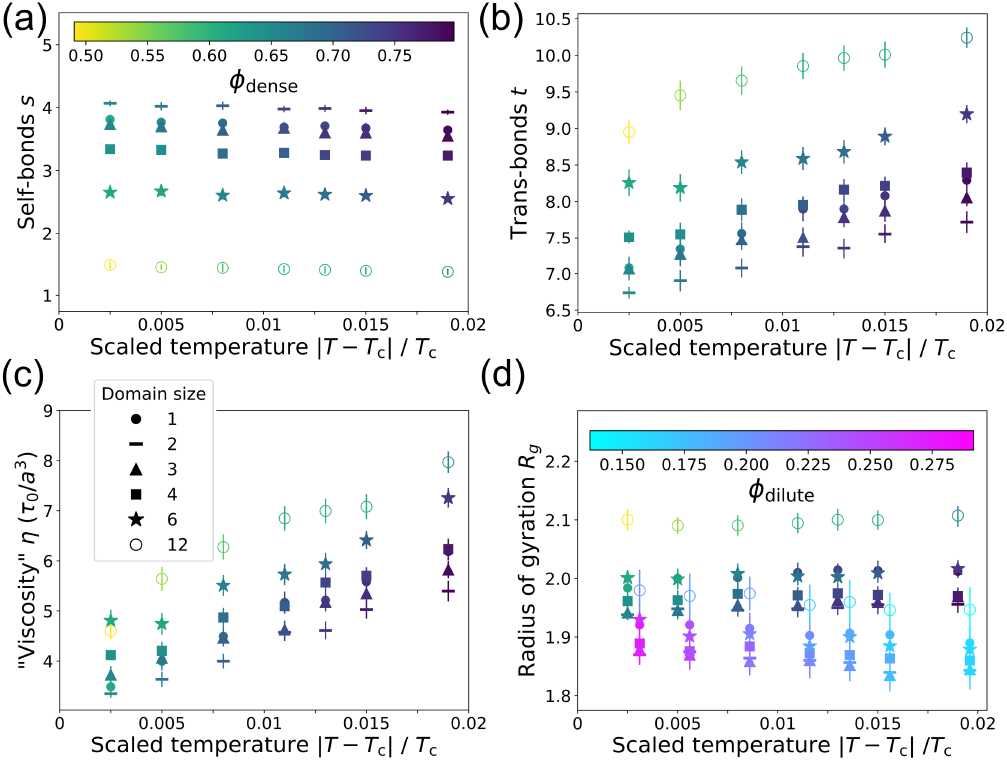
The structure of the dense phase depends on the motif sequence. (a) Number of self-bonds *s* in the dense phase as a function of reduced temperature for domain sequences (symbols as in (c)). Each point shows *s* (mean ± SD) over all configurations with |*ϕ* - *ϕ*_dense_| ≤ 0.01. Color bar: droplet density. (b) Number of trans-bonds *t* (bonds with other polymers) versus temperature as in (a). (c) “Viscosity” (Eq. 5) of the dense phase, shown as in (a). Symbol key applies to all panels. (d) Radius of gyration *R*_g_ of polymers in the dense phase (shown as in (a)) and in the dilute phase. Dilute-phase points show *R*_g_ (mean ± SD) over all configurations with |*ϕ - ϕ*_dilute_| ≤ 0.01. They share reduced temperatures with the dense phase points but are shifted for clarity. Color bar: dilute phase density.

These structural differenceswill affect the physical properties of the dense phase. The timescales of a droplet’s internal dynamics will determine whether it behaves more like a solid or a liquid. We might expect denser droplets to have slower dynamics, so the *ℓ* = 1 and *ℓ* = 2 sequences would be more solid-like. However, the extra inter-polymer bonds at large *ℓ* will slow the dynamics. To disentangle these effects, we estimate the viscosity and polymer-diffusivity by modeling the dense phase as a viscoelastic polymer melt with reversible cross-links formed by trans-bonds. Then the viscosity is expected to scale as (***Rubinstein and Semenov, 2001***)

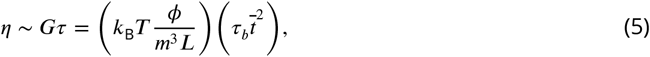

where *G* is the elastic modulus, *τ* is the relaxation time of the polymer melt, and *m* is the monomer length. *τ* depends on the trans-bonds per polymer 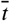 and the bond lifetime *τ_b_* = *τ*_0_ exp(*βε*), where *τ*_0_ is a microscopic time which we take to be sequence-independent. Figure 4(c) shows the dense-phase viscosity calculated using in Eq. 5 the 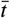 and *ϕ*_dense_ obtained from simulation. We find that sequences with large domains have more viscous droplets due to the strong dependence on inter-polymer bonds, in spite of their substantially lower droplet density. By the same arguments leading to Eq. 5, diffusivity scales as 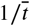, so polymers with large domains will also diffuse more slowly within droplets (Appendix 4, Fig. 13). Thus trans-bonds are the main repository of elastic “memory” in the droplet.

The motif sequence also affects the polymer radius of gyration in both phases (Fig. 4(d)). In the dense phase, polymers with large domains adopt expanded conformations which allow them to form more trans-bonds. Polymers of all sequences are more compact in the dilute phase, where there are fewer trans-bonds and nonspecific interactions with neighbors. Thus self-bonds cause polymers to contract, while trans-bonds cause them to expand.

## Discussion

In summary, we developed a simple lattice-polymer model to study how the sequence of specific-interaction motifs affects phase separation. We found that motif sequence determines the size of the two-phase region by setting the relative entropy of intra-versus inter-molecular bonds. In particular, large domains of a single motif disfavor self-bonds and thus favor phase separation. This is consistent with recent experimental (***Pak et al., 2016***) and theoretical (***Lin et al., 2016; McCarty et al., 2019***) studies on coacervation (phase separation driven by electrostatics) where small charge-domains lead to screening of the attractive forces driving aggregation. However, electrostatic interactions (generic, longer-range, promiscuous) are qualitatively very different from the interactions in our model (specific, local, saturating). This points to a different underlying mechanism: in the former, sequence primarily influences the electrostatic energy of the dense phase, but in the latter, sequence controls the conformational entropy of the dilute phase. Thus specific interactions provide a distinct physical paradigm for the control of intracellular phase separation. While our dilute phase concentrations are large relative to experimental values due to weak nonspecific interactions and the discrete lattice, we expect these sequence-dependent patterns to be quite general. If anything, the self-bond entropy will be even more important at low *ϕ*_dilute_.

We then analyzed how sequence influences condensates’ physical properties such as viscosity and diffusivity. We found that motif sequence strongly affects both droplet density and interpolymer connectivity, and, in particular, that sequences with large domains form more viscous droplets with slower internal diffusion. All sequences expand in the dense phase to form more trans-bonds, and small-domain sequences are the most compact. This contrasts with results for single polyampholyte chains, where “blocky” sequences with large domains are more compact(***Das and Pappu, 2013; Sawle and Ghosh, 2015***). The difference arises because our system includes many polymers interacting with each other and because hairpins are less favored by specific bonds than by longer-range electrostatic interactions.

Taken together, these results suggest that motif sequence provides cells with a means to tune the formation and properties of intracellular condensates. For example, motif stoichiometry could be an active regulatory target - a cell could dissolve droplets by removing just a few binding motifs per polymer through post-translational modifications. The negative correlation between *T*_c_ and *ϕ*_c_ provides another regulatory knob: if a particular condensate density is required at fixed temperature, this can be achieved by either tuning the binding strength or modifying the sequence. However, the physics also implies biological constraints: the same trans-bonds that drive condensation for high-*T*_c_ sequences also lead to high viscosity, which may not be functionally favorable. Key predictions of our model may be tested experimentally using synthetic biopolymers with interaction motifs arranged in domains of different sizes (e.g. using the SIM-SUMO or SH3-PRM systems), then quantifying the relationship between domain size, *T*_c_ or *ϕ*_dilute_, or viscosity/diffusivity.

We have used a simple model of biological condensates to show how the sequence of specificinteraction motifs affects phase separation, thus linking the microscopic details of molecular components to the emergent properties relevant for biological function. What lessons are likely to generalize beyond the details of the model? When nonspecific interactions dominate, forming a dense droplet has a large energetic payoff. When interactions are specific and saturating, however, the energy change is limited and the conformational entropy is expected to play a bigger role. For example, in two-component systems the conformational entropy of small oligimers can stabilize the dilute phase (***Xu et al., 2020; Zhang et al., in press***). Here, we have shown that the conformational entropy of self-interactions can play a similar role, and we use the density of states *g*(*s*) to connect sequence and entropy. Can this framework be extended to other molecular architectures where specific self-interactions are important? For example, mRNA secondary structure can control whether a transcript remains in the dilute phase or enters a protein condensate (***Langdon etal., 2018***). RNA self-interactions could also drive aggregation in disease. Transcripts with nucleotide repeats phase separate more readily than scrambled sequences (***Jain and Vale, 2017***), and it will be interesting to ask how this relates to the robust phase separation of large-domain sequences in the present work. Understanding the general role of the entropy of self-interactions will prove useful if it allows us to gain insight into biomolecular phase separation by simply analyzing the properties of single molecules or small oligomers rather than necessarily tackling the full many-body problem. Many open questions remain, however, and we hope our work encourages further research across a range of theoretical and experimental systems.

## Acknowledgments

We thank O. Kimchi, E. King, and J. Steinberg for valuable conversations about RNA phase separation. This work was supported in part by the National Science Foundation, through Center for the Physics of Biological Function Grant PHY-1734030 (to B.G.W.), and National Institutes of Health Grant R01 GM140032 (to N.S.W.).

## Methods and Materials

We performed Monte Carlo simulations in the Grand Canonical Ensemble on a 30×30×30 FCC lattice, corresponding to a volume of *V* = 30^3^ lattice sites, with periodic boundary conditions. When “A” and “B” monomers occupy the same site, they form a bond with energy *ε*. Other overlaps are forbidden. When two monomers of any type occupy adjacent lattice sites, they have an attractive nonspecific interaction energy *J*. Thus each lattice site *i* has a bond occupancy *q_i_* ∈ [0,1] and a motif occupancy *r_i_* ∈ [0,1,2]. The Hamiltonian for our system is therefore

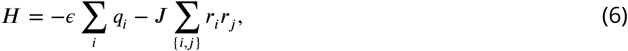

where the brackets indicate summation over adjacent lattice sites. Each simulation has fixed control variables *β* = 1/*k_B_T* and polymer chemical potential *μ*. We use simulated annealing to cool the system to the final temperature, and after reaching that temperature to ensure the system has thermalized we only use data from the last 80% of steps. The total number of Monte Carlo steps varies, but is around 4.5 · 10^8^ for critical point simulations. In each Monte Carlo step, we update the system configuration by proposing a move from the move-set defined in Fig. 5. Moves (a-c) are standard polymer moves. We include contraction and expansion moves (Fig. 5(d) and (e)) which allow contiguous motifs to form and break bonds. The FCC lattice has coordination number *z* = 12, so there are 12 states that can transition into any one contracted state. Thus it is necessary to propose expansions at 12 times the rate of contractions to satisfy detailed balance. We also allow clusters of polymers connected by A-B overlap to translate by one site so long as no overlap bonds are formed or broken.

**Figure 5.**
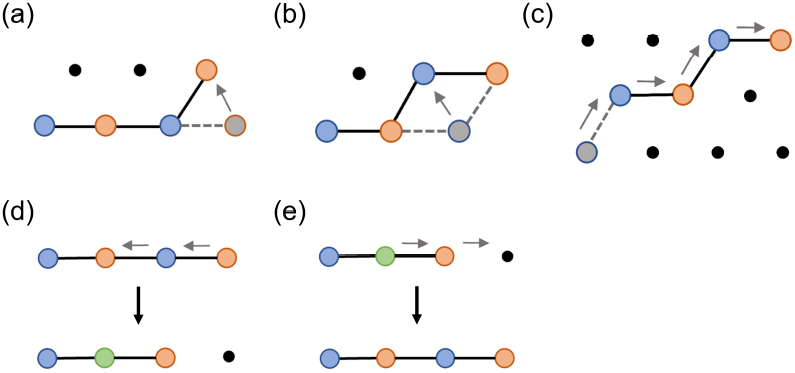
The polymer moves used to update Monte Carlo simulations at each step. We also allow translation of connected clusters of polymers and insertion/deletion of polymers. (a) End move. (b) Corner move. (c) Reptation. (d) Contraction. (e) Expansion.

To include insertions and deletions of polymers, we assume the existence of a reservoir of polymers of chemical potential *μ*, which we can adjust. Because inserting a polymer tends to increase the configurational entropy of the system, we adopt the common convention of shifting *μ* by the entropy of an ideal polymer: *μ* ≡ *μ*_0_ + ln(*z* + 1)^*L*-1^, where the “+1” in *z* +1 comes from allowing the “walk” to remain on the same site and form a contiguous bond (see Fig. 5(d)-(e)). We then remove the shift with a prefactor in the acceptance probabilities (Eq. 12). This convention allows us to simulate the dilute phase without setting *μ* to a large negative value.

In our Monte Carlo move set, we allow for the deletion of any polymer, and require that insertion moves satisfy detailed balance with respect to deletions. This still allows for considerable freedom in the insertion algorithm. Naively, we might insert polymers as random walks, but for a dense system most such random walks will be disallowed because of forbidden overlaps. For efficiency, we therefore implemented a form of Configurational-Bias Monte Carlo (*CBMC*)(***Frenkel and Smit, 2002***). Specifically, we insert the head of a polymer at a randomly chosen site, and then perform a biased walk along an allowed path, keeping track of the number of available choices at each step to generate a “Rosenbluth weight” *R*:

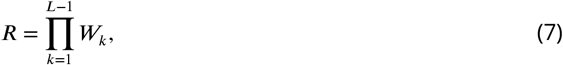

where *W_k_* is the number of allowed sites for monomer *k* +1 starting from the position of monomer *k*. The probability of this insertion move is therefore

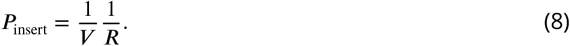

The CBMC algorithm satisfies detailed balance so long as the net flow of probability between any two configurations *x*_1_ and *x*_2_ is zero. In words, this imposes the condition

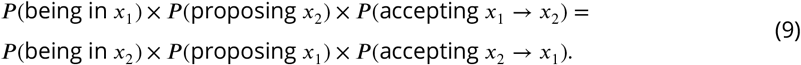

In our system, if configuration *x*_1_ has polymer number *N* and energy *E_N_* and *x*_2_ has polymer number *N* + 1 and energy *E_N_*_+1_, Eq. 9 becomes

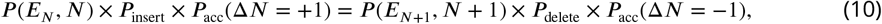

where *P*(*E,N*) = exp(-*βE* + *βμN*)/*Z* is the equilibrium probability of the state. CBMC leads to the *P*_insert_ in Eq. 8. *P*_delete_ = 1/(*N* + 1), because polymers are chosen randomly for deletion. This leads to the following condition on the acceptance probabilities:

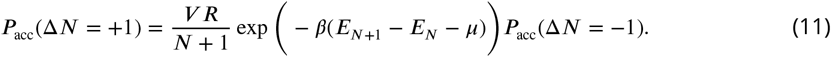

The acceptance probabilities given below in Eq. 12 satisfy this condition and also incorporate the multicanonical sampling described next.

We determine the phase diagram using histogram reweighting (Panagiotopoulos et al., 1998) of *P*(*N,E*), where *N* is the polymer number and *E* is the total system energy. This allows us to extrapolate a histogram *P*(*N, E*) obtained at *β*_0_, *μ*_0_ to 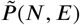 at nearby *β*_1_,*μ*_1_. First we determine the approximate location of the critical point, then run a sufficiently long simulation to obtain a converged *P* (*N,E*). We determine the exact location of the critical point by finding the *β*_c_,*μ*_c_ where 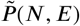 matches the universal distribution known for the 3D Ising model (Tsypin and Blöte, 2000). (Because polymer models lack the symmetry of the Ising model, we also must fit a “mixing parameter” *x* which determines the order parameter *N-xE* (***Wilding, 1997***).) In principle, we could find the binodal at temperature *T* < *T*_c_ (*β* > *β*_c_) by determining *P_β_*(*N, E*), then reweighting the histogram to the *μ*\* at which *P_β_*(*N*) has two peaks with equal weight. The phase boundaries *ϕ*_dilute_ and *ϕ*_dense_ would then be the means of these peaks, which we could find by fitting *P_β_* (*N*) to a Gaussian mixture model. However, determining the relative equilibrium weights of the two phases requires observing many transition events, which are very rare at temperatures substantially below *T*_c_. To circumvent this difficulty, we use multicanonical sampling (***Wilding, 1997***): Once we have *P_βc_* (*N,E*) at the critical point, we use reweighting to estimate 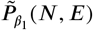 at a slightly lower temperature *β*_1_. When we perform the new simulation at *β*_1_, we use a modified Hamiltonian 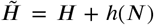, where 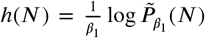. (Note that *h*(*N*) is only defined over the range of *N* between the two peaks.) This yields 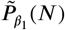, which is unimodal and flat-topped with respect to *N* rather than bimodal, and thus allows rapid sampling of the full range of relevant values of *N*. Figure 6(a) shows an example distribution 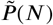. Finally, we use reweighting to remove *h*(*N*) and study the true histogram *P_β1_*(*N,E*), as in Fig. 6(b). We apply this procedure iteratively to obtain the phase boundary at lower and lower temperatures. Combining multicanonical sampling with Configurational-Bias Monte Carlo, our acceptance probabilities become

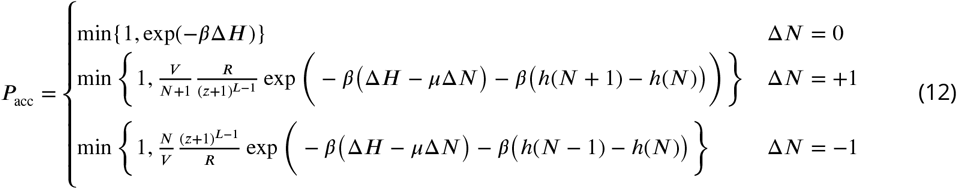

*Single-polymer properties*. The density of states *g*(*s*) is the number of configurations of an isolated polymer with *s* self-bonds. We extract *g*(*s*) by performing Monte Carlo simulations of the polymer over a range of *β* values. The distributions are then combined using the multihistogram method, and inverted to determine the density of states (***Landau and Binder, 2014***).

**Figure 6.**
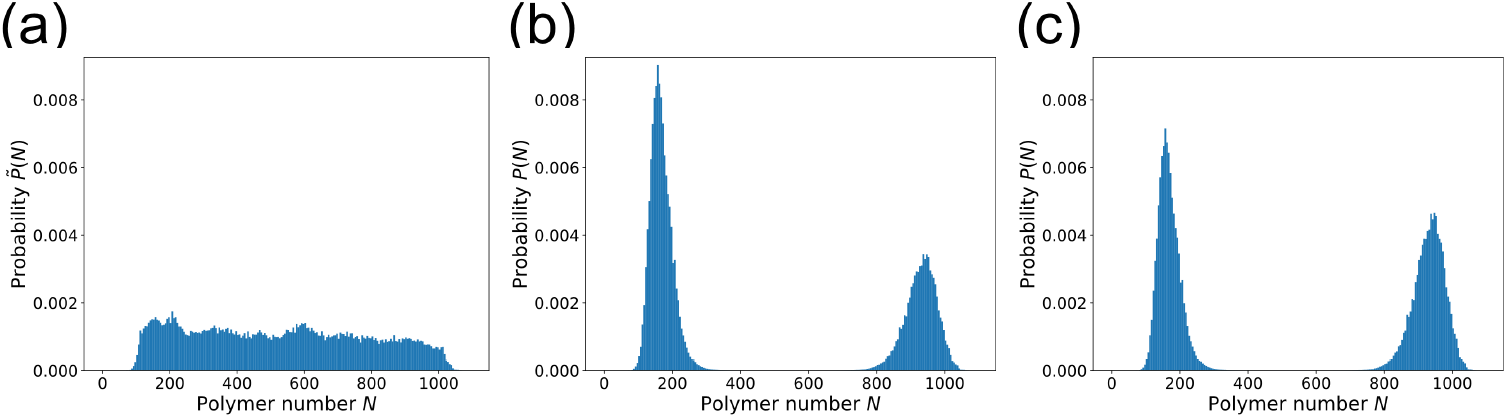
Multicanonical sampling makes it possible to determine the phase boundary at temperatures substantially below *T*_c_. (a) The polymer number distribution 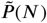 produced in a multicanonical simulation with 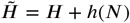. Domain sequence with *ℓ* = 2, *βε* ≈ 0.94, *J* = 0.05*ε*. (b) The true distribution *P*(*N*), obtained by reweighting 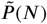 from (a) to remove *h*(*N*). (c) The distribution at the phase boundary, obtained by reweighting (b) to the chemical potential *μ*\* at which both peaks have equal weight.

## Appendix 1 Mean-field theory

We aim to find the partition function *Z* for a system with *N* identical, interacting polymers on a lattice with *V* sites. Each polymer has *a* A motifs, *b* B motifs, and length *L* = *a* + *b*. We label the state of polymer *i* by the number of self-bonds *s_i_* and trans-bonds *t_i_*. Then the total number of self-bonds is *S* ≡∑_i_ *s_i_*, and the total number of trans-bonds is 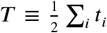. In our approach, each polymer forms self-bonds according to its own full degrees of freedom encoded in the density of states *g*(*s*). However, we approximate the inter-polymer interactions within a mean-field approach. The full partition function for our system is then given by

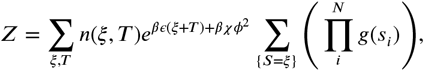

where *n*(ξ,*T*) is the combinatorial term for counting states with *T* A-B overlap bonds (given total self-bonds) and the second sum is over all configurations where *S* = ξ. The parameter *χ* quantifies the strength of two-body nonspecific interactions, e.g. as appears in Flory-Huggins theory. We make the approximation that in the thermodynamic limit, *Z* is dominated by the largest term:

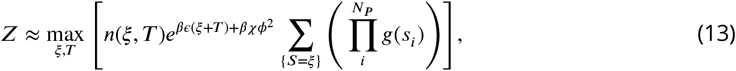

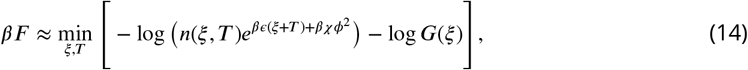

where *G*(ξ) is the entropy associated with forming *S* = ξ self-bonds.

First we calculate *n*(ξ,*T*) = *n*_steric_ × *n*_trans_. *n*_steric_ is the number of allowed ways to place the polymers on the lattice and *n*_trans_ is the number of ways to form *T* trans-bonds. To find *n*_steric’_, we ignore chain connectivity and simply count the number of ways of choosing *N*〈*l*〉 sites on a lattice with *V* sites, where

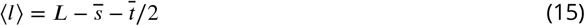

is the mean number of sites occupied by a polymer. We account for excluded volume using a semi-dilute approximation that the probability of placing monomer *k* successfully is the fraction of empty sites remaining:

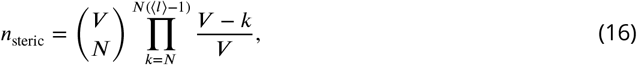

where 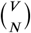 counts the center-of-mass, or equivalently “polymer head,” degrees of freedom. We find *n*_trans_ by assuming that each protein sees the others as a mean-field cloud of motifs with which it can form A-B overlap bonds depending on the overall motif density. Then

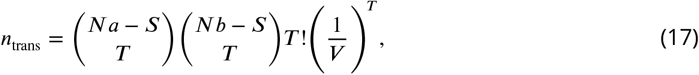

where the first two terms count the number of ways to choose *T* A motifs and *T* B motifs, given that *S* of each are already in self-bonds. *T*! is the number of ways to pair the chosen motifs, and the final term is the mean-field probability that two motifs are close enough to form a bond. (This is simply an extension of Semenov and Rubinstein’s sticker model to two sticker types on a lattice (***Semenov and Rubinstein, 1998***).)

Nowwe calculate *F_G_* (*ξ*) ≡ -log *G*(*ξ*), the entropy of having exactly *S* = *ξ* self-bonds. The difficulty arises from the restricted sum: we only want to count states with the correct total number of self-bonds. However, we can relax this restriction and require instead that 〈*S*〉 = *ξ*. Formally, this is equivalent to working in a “Grand Canonical Ensemble” for self-bonds, where a reservoir imposes a chemical potential *w*. In the thermodynamic limit, fluctuations vanish and all ensembles yield equivalent macrostates. Thus we can calculate *β*Ω = - log *Z*_gc_ (where Ω is the grand potential and *Z*_gc_ the grand canonical partition function), and use the Legendre transform *F_G_*(*ξ*) = Ω + *wξ*/*β*.

Calculating *Z*_gc_ is relatively straightforward:

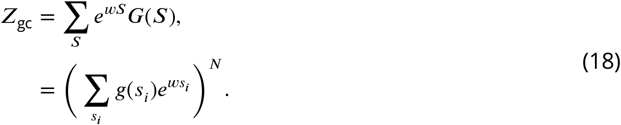

Then *w* = *w*(*ξ*) is fixed by requiring that 〈*S*〉 = ξ. Recall that 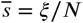, so

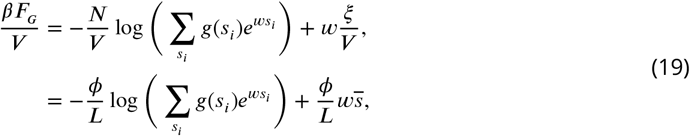

where *ϕ* is the monomer density *NL*/*V*. Combining this with Eqs. 16 and 17, we obtain the full free-energy density:

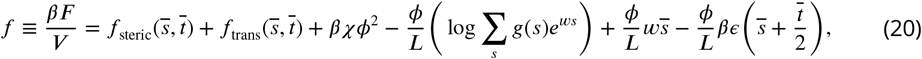

where

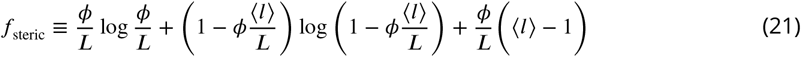

and

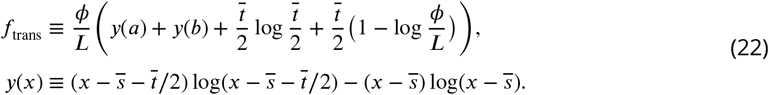

At every *ϕ*, we evaluate Eq. 20 with the average bond values 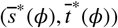 which minimize *f* and the *w* which fixes 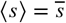. This yields *f*(*ϕ*) which we use to calculate the binodal and spinodal curves.

Regarding the nonspecific interaction parameter *χ*, density fluctuations make it difficult to map the simulation *J* to *χ*, so we simply use the mean-field relation *χ* = -*VJz*/2, where *z* is the lattice coordination number. This yields theoretical *T*_c_ values which differ numerically from the simulations but accurately reproduce the sequence hierarchy.

## Appendix 2 Dense-phase correlations

From simulations, the *ℓ* = 1 sequence has a *T*_c_ between that of *ℓ* = 3 and *ℓ* = 4, whereas the mean-field theory predicts that *ℓ* = 1 would have a *T*_c_ very close to that for *ℓ* = 2. Why is the *ℓ* = 1 sequence better at phase separating than the mean-field theory predicts? In the theory, sequence only appears in *g*(*s*), the density of states for self-bonds. We thus assume that sequence does not directly affect inter-polymer interactions and that trans-bonds are uncorrelated. However, this assumption neglects the fact that a bond is between two polymers. We can quantify this correlation by looking at trans-bond “segments.” Trans-bonds are considered to be in a segment of length *λ* if two polymers have *λ* trans-bonds, and all involved monomers are contiguous on both polymers (Fig. 7(a) *Inset*). Essentially, trans-bond segments form when two polymers are lying on top of each other. Figure 7(a) shows the probability that each trans-bond is in a segment of length *λ* in an NVT simulation with *ϕ* = 0.3. For all sequences, the most probable segment length is 1. However, *ℓ* = 1 and *ℓ* = 12 both have relatively high probabilities of forming longer segments (these two curves overlap). As a result of these correlations, the dense phase is more favorable for these sequences than is predicted by the theory, and this leads to their higher *T*_c_ values.

**Figure 7.**
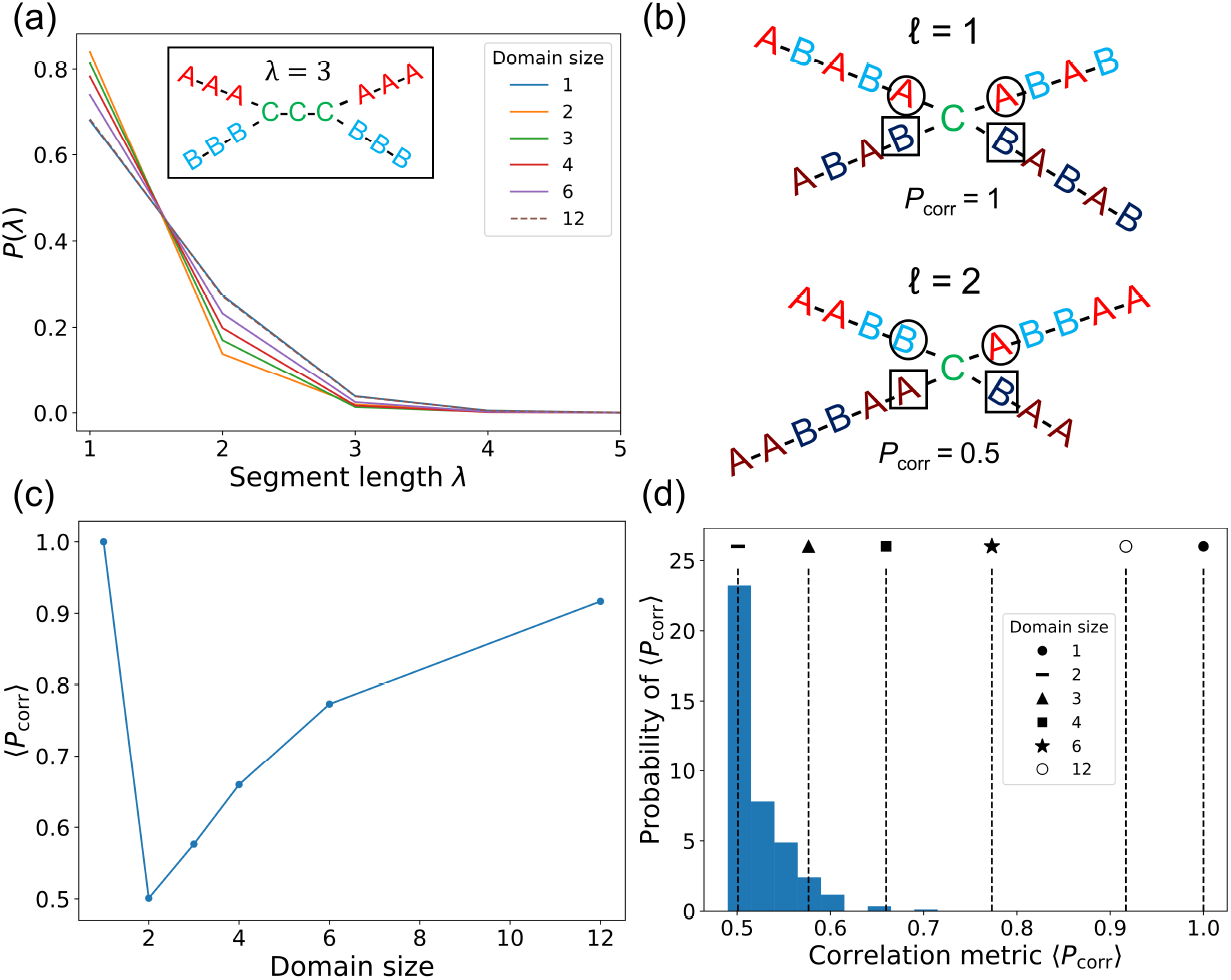
The *ℓ* = 1 polymer has correlated trans-bonds in the dense phase. (a) Probability that a trans-bond is in a segment of length *λ*, meaning it has *λ* bonds with the same partner, and all *λ* monomers are contiguous on both polymers. Data from snapshots of an NVT simulation with *ϕ* = 0.3, *βε* = 1.25, *J* = 0.05*ε*. *Inset*: A trans-bond segment of *λ* = 3, between one polymer with (*a, b*) = (9,0) and another polymer with (*a, b*) = (0,9). (b) Example *P*_corr_ for bonds between *ℓ* = 1 (*top*) and *ℓ* = 2 (*bottom*) polymers. Motifs from polymer 1 and 2 are distinguished by lighter and darker shades, respectively. Bond-adjacent monomers are marked by circles for polymer 1 and squares for polymer 2. The pictured bond’s *P*_corr_ is the fraction of square-circle pairs that are A-B. (c) Trans-bond correlation probability 〈*P*_corr_〉 for domain sequences, where the brackets denote averaging over initial bonds. (d) Distribution of 〈*P*_corr_〉 for 20,000 scrambled sequences with *a* = *b* =12. Values for the domain sequences are marked.

We can quantify a sequence’s tendency to form correlated segment bonds by defining a correlation probability *P*_corr_. Consider two polymers which form a bond between monomers *i* and *j*. Now pair up neighboring monomers: the four unique possibilities are (*i* −1, *j* −1), (*i* −1, *j* + 1), (*i* + 1, *j* −1), and (*i* + 1, *j* + 1). *P*_corr_ is the probability that these monomers will form a valid A-B bond instead of an invalid overlap. Figure 7(b) shows examples for *ℓ* = 1 and *ℓ* = 2 sequences. Every possible initial bond (*i,j*) has its own *P*_corr_, and so we average this *P*_corr_ over all possible bonds. This yields 〈*P*_corr_〉, a sequence-specific metric for trans-bond correlations. Figure 7(c) shows 〈*P*_corr_〉 for the domain sequences, and we observe that it is monotonic in domain size *except* for *ℓ* = 1, which has a 〈*P*_corr_〉 similar to *ℓ* = 12. This explains why these two sequences have similar segment probabilities in Fig. 7(a), and why *ℓ* = 1 is better at phase separating than expected from *g*(*s*) alone. In Appendix 3 below, we incorporate 〈*P*_corr_〉 into a “condensation parameter” that successfully predicts the *T*_c_ hierarchy observed in simulation. Figure 7(d) shows the distribution of 〈*P*_corr_〉 for 20,000 random sequences with *a* = *b* = 12. The distribution is strongly peaked at low values, comparable to the *ℓ* = 2 sequence. This suggests that the *ℓ* = 1 and *ℓ* = 12 domain sequences are atypical in their tendency to form correlated trans-bonds, so the mean-field theory that neglects these correlations should perform well for generic sequences.

## Appendix 3 Condensation parameter Ψ

Although our mean-field theory does a good job explaining sequence-driven patterns in *T*_c_, it would be convenient to have an order parameter that is simpler to compute but that retains some of the same predictive power. According to our results, such a metric should take into account the density of states *g*(*s*), the motif stoichiometry *a, b,* and the correlation metric 〈*P*_corr_〉. Thus we propose as a metric the condation parameter Ψ:

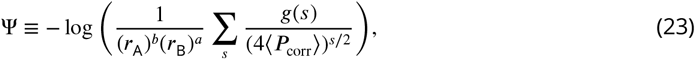

where the motif ratios are given by *r*_A_ = *a/L* and *r*_B_ = *b/L*. The role of *g*(*s*) is intuitive: the easier it is to form self-bonds, the less a polymer will tend to condense. The factor 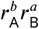 characterizes the probability of placing *a* A motifs and *b* B motifs in the dense phase without disallowed overlap. (The mean-field motif placement probability depends on the density *ϕ*, but this effect is not sequence-dependent.) Finally, we normalize *g*(*s*) by the tendency to form correlated trans-bonds in the dense phase. This tendency enhances the favorability of the dense phase, and we quantify it with 〈*P*_corr_〉. The factor of 1/2 in *s*/2 is due to the fact that two trans-bonds/polymer are required to lower the energy by *ε*/polymer, and the factor of 4 is the number of pairs of bond-adjacent monomers (Fig. 7(b)). Although this metric is only heuristic, it successfully captures the *T*_c_ patterns without multi-polymer simulations (Fig. 3(c)).

One limitation of the condensation parameter is that it still requires knowledge of *g*(*s*) for each sequence. Is it possible to characterize the tendency of a sequence to phase separate without any simulations? In Fig. Fig. 3(c) of the main text, we replace ∑_*s*_ *g*(*s*) with a theoretical calculation of *g*(1)/*g*(0) that uses established scaling relations for the number of self-avoiding walks and the number of self-avoiding loops ***De Gennes, 1979***). This gives

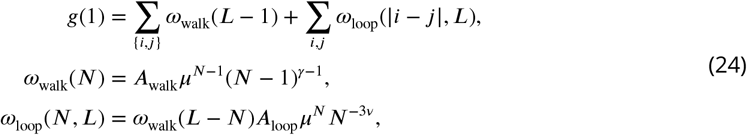

where *ω*_walk_(*L* −1) is the number of self-avoiding walks when a polymer of length *L* forms a contiguous bond (shortening it by 1 monomer), and *ω*_loop_(*N, L*) counts the number of self-avoiding loops of length *N*. We model the entropy of the polymer outside the loop as a self-avoiding walk of length *L-N*. The sums are over all possible contiguous bonds and loops, which depend on the compatibility of motifs *i* and *j*. The exponents *γ* = 1.157 and *v* = 0.588 are universal, and *μ* = 10.037 on the FCC lattice (this coefficient *μ*, which is standard notation, is not to be confused with the chemical potential *μ* in our simulations). The scaling amplitudes *A*_walk_ and *A*_loop_ are not universal, so we determine their relative magnitude by fitting to *g*(1) from the Monte Carlo *g*(*s*) for a single sequence. With this one fitting parameter, we can rapidly evaluate Ψ for new sequences with no additional simulations or calculations. Specifically, we perform a linear fit of Ψ to *T*_c_ for the domain sequences (Fig. 8) and obtain *T*_c_ for any new sequence from its Ψ value. This procedure allows us to generate the *T*_c_ distribution in Fig. 3(d) in seconds. A Python script to calculate Ψ and *T*_c_ for arbitrary sequences is available at https://github.com/BenjaminWeiner/motif-sequence/tree/master/condensation%20analysis.

**Figure 8.**
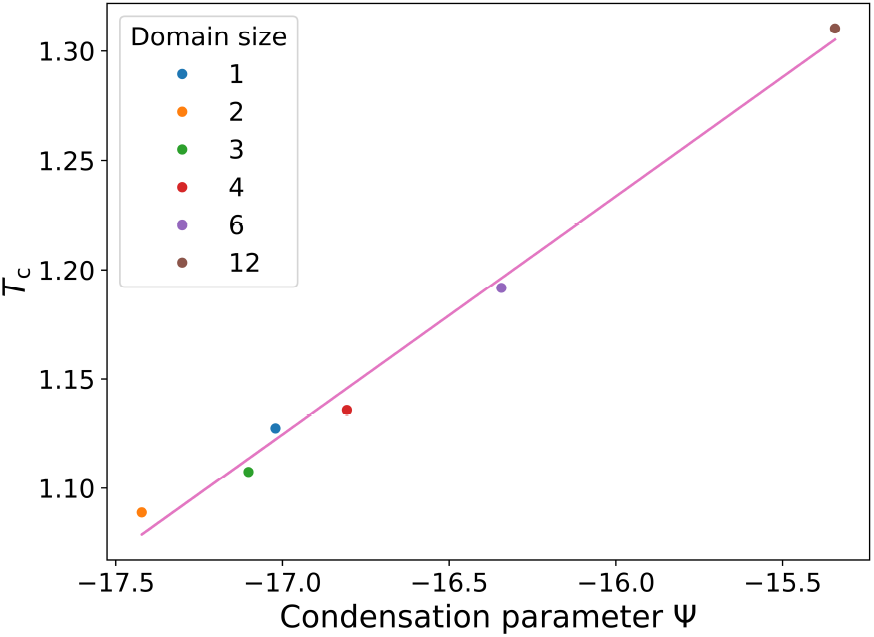
The linear fit between *T*_c_ from Monte Carlo simulations and Ψ calculated via Eq. 24. Slope=0.1089, intercept=2.9767.

## Appendix 4 Additional figures

**Figure 9.**
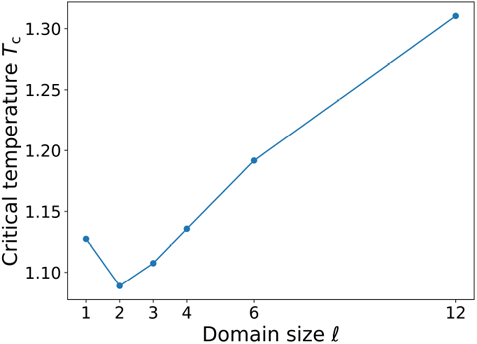
The critical temperatures of *L* = 24 domain sequences. *T*_c_ is monotonic in domain size *ℓ* except for the *ℓ* = 1 sequence, which has strong trans-bond correlations (see Appendix 2). Mean ± SD over three replicates. (Temperature uncertainties are too small to see.)

**Figure 10.**
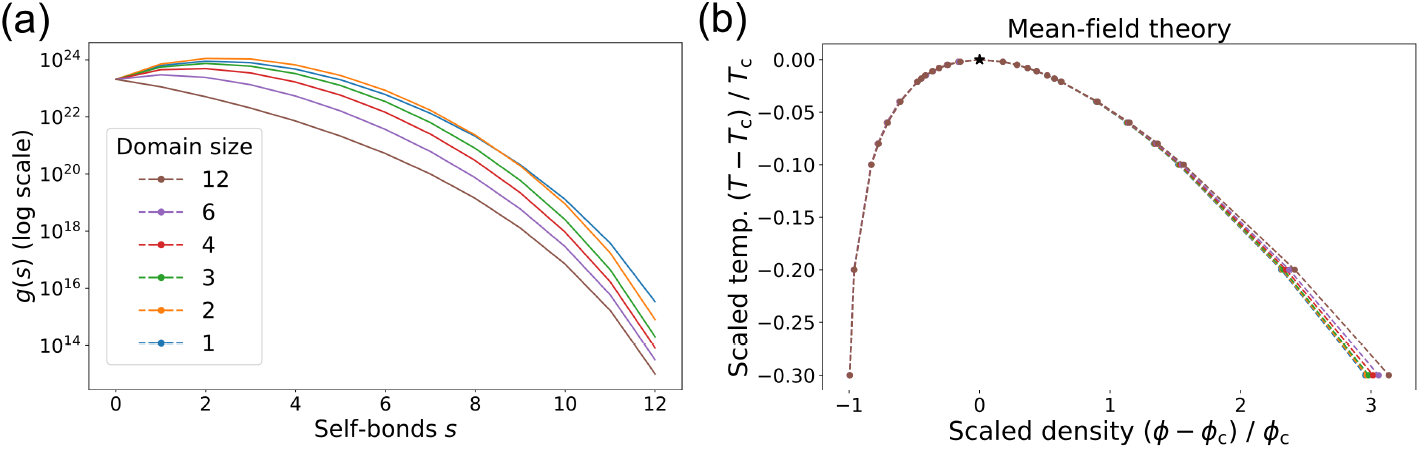
(a) The density of states *g*(*s*), i.e. the number of ways a given sequence can form *s* bonds with itself, semi-log plot. Domain sequences have large differences in *g*(*s*) even for relatively rare states with large *s*. Domain color code applies to all panels. (b) The phase diagram from the mean-field theory, rescaled by the critical temperature *T*_c_ and critical density *ϕ*_c_.

**Figure 11.**
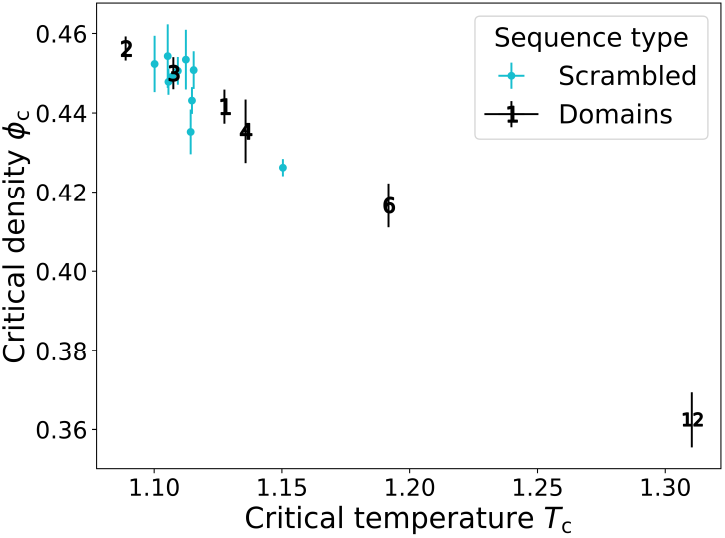
Critical temperatures and critical densities of *L* = 24 domain sequences and scrambled sequences, all with *a* = *b* =12. For the domain sequences, the plot markers denote domain size *ℓ*. Scrambled sequences cluster around the *ℓ* = 3 domain sequence, motivating the use of this sequence as the starting point for stoichiometry mutations in Fig. 3(b). Mean ± SD over three replicates. (Temperature uncertainties are too small to see.)

**Figure 12.**
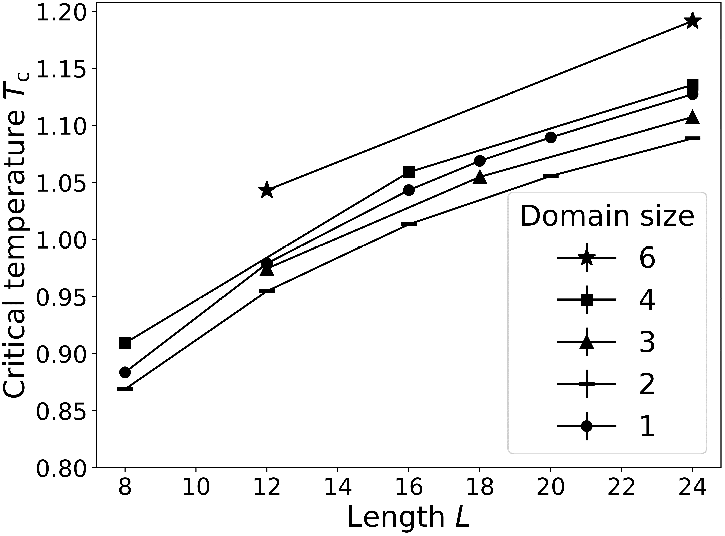
*T*_c_ as a function of length for sequences with different domain sizes. mean ± SD over three replicates. (temperature uncertainties are too small to see.) the *T*_c_ hierachy is preserved across sequence lengths. thus domain size is a robuts predictor of phase separation via its relationship with self-bond entropy.

**Figure 13.**
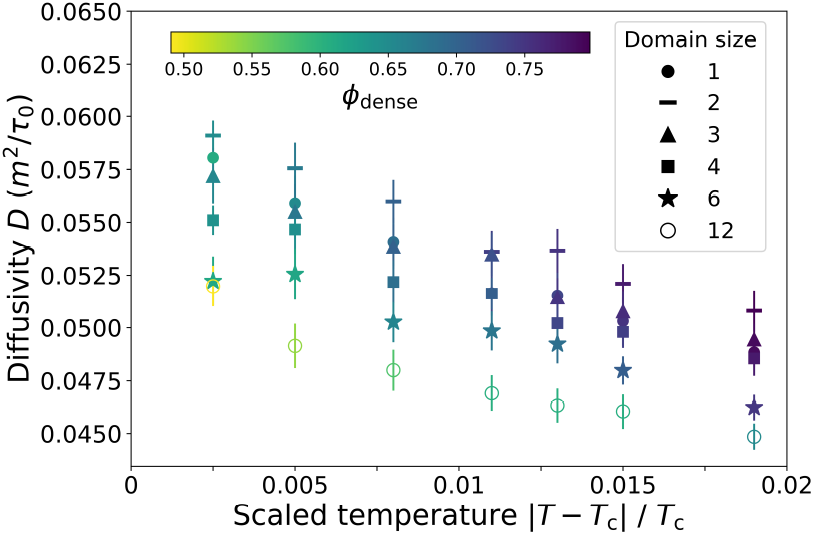
Using “sticky rouse model” for unentangled polymer dynamics in a melt with cross-links (Rubinstein and Semenov, 2001), the dense-phase diffusivity 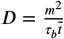, where *m* is the monomer size and *τ_b_* = *τ*_0_ exp(*βε*) is the bond lifetime is plotted as a function of scaled temperature. for all sequence, lower temperatures correspond to higher densities and slower polymer diffusion. importantly, the sequence with large domain sizes and many trans-bonds (e.g. *ℓ* = 12 and *ℓ* = 6) have smaller *D,* in spite of their lower density. this coincides with the viscosity results fig. 4 of the main text, where the trans-bonds dominate the physical properties of the droplet. color bar: droplet density.

